# Combined absence of TRP53 target genes ZMAT3, PUMA and p21 cause a high incidence of cancer in mice

**DOI:** 10.1101/2022.09.06.506741

**Authors:** Margs S. Brennan, Kerstin Brinkmann, Geraldine Healey, Lahiru Gangoda, Andreas Strasser, Marco J. Herold, Ana Janic

## Abstract

Transcriptional activation of target genes is essential for TP53-mediated tumour suppression, though the roles of many TP53-activated target genes in tumour suppression remains poorly understood. Knockdown of ZMAT3 in haematopoietic stem/progenitor cells by shRNA caused leukaemia only with the concomitant absence of the pro-apoptotic BCL-2 family member PUMA and the CDK inhibitor p21. We were interested to further investigate the role of ZMAT3 in tumour suppression beyond the haematopoietic system. Therefore, we generated *Zmat3* knockout and compound gene knockout mice, lacking *Zmat3* and *p21*, *Zmat3* and *Puma* or all three genes. *Puma^−/−^p21^−/−^Zmat3^−/−^* triple knockout mice developed tumours at a significantly higher frequency compared to wild type, *Puma^−/−^Zmat3^−/−^* or *p21^−/−^ Zmat3^−/−^*deficient mice. Interestingly, we observed that the triple and *Puma^−/−^Zmat3^−/−^* double deficient animals succumbed to lymphoma, while *p21^−/−^Zmat3^−/−^* animals developed mainly solid cancers. This analysis suggests that in addition to ZMAT3 loss, additional TRP53-regulated processes must be disabled simultaneously for TRP53-mediated tumour suppression to fail. Interestingly, our findings also reveal that the absence of different TRP53 regulated tumour suppressive processes changes the tumour spectrum, which suggests that different TRP53 tumour suppressive pathways are more critical in different tissues.

## INTRODUCTION

The tumour suppressor TP53 (mouse TRP53, often referred to as p53) is frequently mutated gene in human cancer (Aubrey *et al*, 2018; Kastenhuber & Lowe, 2017; Vousden & Lane, 2007). Although TP53-mediated transcriptional regulation of diverse cellular responses is known to be critical for its ability to prevent the development of cancer, the role of many TP53 target genes in tumour suppression remains unclear (Kastenhuber & Lowe, 2017; Boutelle & Attardi, 2021). Recent studies in mice sought to identify the mechanisms that are critical for TRP53-mediated tumour suppression, using sensitised shRNA gene knock-down and CRISPR/Cas9 knock-out screens in cells in culture and *in vivo* (Janic *et al*, 2018; Bieging-Rolett *et al*, 2020). Amongst the hits identified in these screens, the RNA binding protein ZMAT3 (Bersani *et al*, 2014; Vilborg *et al*, 2011) was found to be a potent tumour suppressor. ZMAT3 (also known as WIG-1) is an RNA-binding zinc-finger protein that is involved in regulating alternative splicing and as such is expressed in a broad range of tissues (Bieging-Rolett *et al*, 2020). ZMAT3 was reported to control splicing of mRNAs, including those encoding the TRP53 regulators, MDM2 and MDM4, several other splicing factors (e.g. HNRNPDL, DHX9) and the cell adhesion and stem cell marker CD44 (Bieging-Rolett *et al*, 2020; Muys *et al*, 2020). ZMAT3 appears to be under the direct transcriptional control of TRP53 in various cell types (Janic *et al*, 2018; Bieging-Rolett *et al*, 2020; Vilborg *et al*, 2011; Brady *et al*, 2011b), indicating its broad role for tumour suppression in diverse tissues. The absence of TRP53 causes a reduction in *Zmat3* expression in a range of human carcinomas (e.g. breast, lung) and high levels of *Zmat3* expression in malignant cells predict increased patient survival in certain cancers (Bieging-Rolett *et al*, 2020). Moreover, an analysis of the Project Achilles CRISPR/Cas9 gene knock-out screening data in human cancer cells revealed that ZMAT3 inactivation enhances proliferation of malignant cells with wt TP53, further supporting a role for ZMAT3 in tumour suppression acting downstream of TP53 (Bieging-Rolett *et al*, 2020). Of note, CRISPR/Cas9-mediated loss of ZMAT3 enhanced tumorigenesis in autochthonous mouse models of lung adenocarcinoma and hepatocellular carcinoma (Bieging-Rolett *et al*, 2020). However, unlike loss of TRP53, the absence of ZMAT3 did not have marked impact on the rate of tumour development or severity of malignant disease in the context of murine c-MYC-driven lymphomagenesis or mutant *Kras^G12D^*-driven lung adenocarcinoma development (Best *et al*, 2020). This indicates that the relative importance of ZMAT3 in TRP53-mediated tumour suppression may vary depending on cell type and/or the oncogenic drivers present in the emerging neoplastic cell population. *In vivo* studies have shown that shRNA-mediated silencing of ZMAT3 in haematopoietic stem/progenitor cells (HSPCs) caused development of leukaemia/lymphoma in transplant recipient mice only when PUMA and p21, the critical effectors of TRP53-mediated apoptosis (Villunger *et al*, 2003; Jeffers *et al*, 2003) and cell cycle arrest (Deng *et al*, 1995) respectively, were also absent (Janic *et al*, 2018). A limitation of these studies was that ZMAT3, PUMA and p21 were removed only in the haematopoietic compartment. Therefore, it remains unclear what impact the absence of ZMAT3 together with the loss of TRP53 induced apoptosis and cell cycle arrest/cell senescence might have in other tissues. To address this question, we generated mice lacking ZMAT3, PUMA and p21. We found that these triple knockout (TKO) mice spontaneously developed malignancy at a considerably higher frequency compared to wt control as well as *Puma^−/−^Zmat3^−/−^* and *p21^−/−^Zmat3^−/−^* double knockout (DKO) mice. Most of the tumours from the TKO mice were of haematopoietic cell origin. These findings demonstrate that this combination of defects in TRP53-activated cellular responses, even if present in all cells, predominantly causes leukaemia/lymphoma. They also reaffirm that defects in multiple TRP53-activated cellular responses are needed to drive tumorigenesis.

## RESULTS

### Adult pre-neoplastic *Puma^−/−^p21^−/−^Zmat3^−/−^* mice have only minor abnormalities in their haematopoietic cell populations

To investigate the impact of combined loss of ZMAT3, PUMA and p21 in TRP53-mediated tumour suppression we crossed *Zmat3^−/−^* with *Puma^−/−^p21^−/−^* mice to obtain *Puma^−/−^Zmat3^−/−^, p21^−/−^Zmat3^−/−^* and *Puma^−/−^p21^−/−^Zmat3^−/−^* animals (Figure 1A). *Puma^−/−^Zmat3^−/−^*, *p21^−/−^Zmat3^−/−^* and *Puma^−/−^p21^−/−^Zmat3^−/−^* offspring were born at the expected Mendelian ratios of inheritance observed for heterozygous crosses of each allele within the colony (Table S1), as well as from *Puma^+/−^Zmat3^+/−^* and *p21^+/−^Zmat3^+/−^* di-hybrid inter-crosses (Table S2). The DKO and TKO mutant mice reached adulthood without any notable defects.

**Figure 1.**
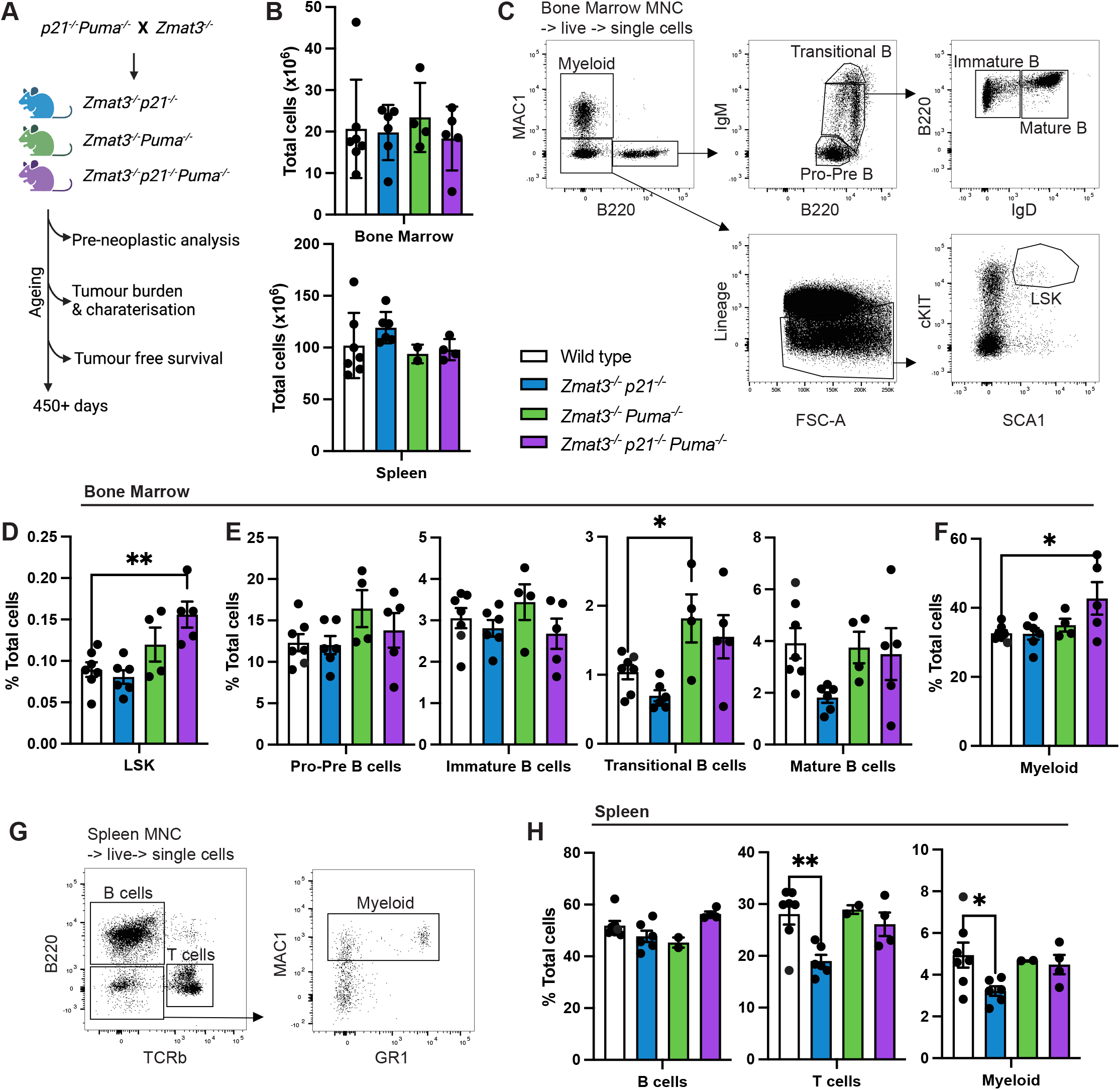
Lymphoid organ analysis of young adult mice of the indicated genotypes. **A)** Schematic for tumour development study. Mouse cohorts were analysed for pre-neoplastic phenotypes and monitored for tumour-free survival. **B-H)** Single-cell suspensions were prepared from spleen, thymus, peripheral blood and bone marrow of *Puma^−/−^p21^−/−^Zmat3^−/−^* (N=5), *Puma^−/−^Zmat3^−/−^* (N=2-4), *p21^−/−^Zmat3^−/−^* (N=6), and wt (N=7) mice and the indicated haematopoietic cell subsets were examined by immunostaining and FACS analysis. **B)** Total cell counts for bone marrow (1 femur, top) and whole spleen (bottom). **C)** Representative FACS plots from wt mice that indicate gating strategy to identify the cell populations of interest in the bone marrow: myeloid (MAC1^+^), LSK (Lineage^−^SCA1^+^cKIT^+^) and B lymphoid cells; pro-pre (B220^lo^IgM^−^), immature (B220^+^IgM^lo^ IgD^−^), transitional (B220^+^IgM^hi^) and mature (B220^+^IgM^med^IgD^+^) B lymphoid cells. The lineage marker antibody cocktail included antibodies against NK1.1, TER119, Ly6G, F4/80, CD2, CD4 and CD8. **D-F)** Percentages of the indicated cell subsets in the bone marrow in mice of the indicated genotypes. **G)** Representative FACS plots of wt mice indicate gating strategy to identify cell populations in the spleen: B cells (B220^+^), T cells (TCRβ^+^) and myeloid cells (Mac1^+^B220^−^ TCRβ^−^). H) Percentages of the indicated cell subsets in the spleen of mice of the indicated genotypes. Data represent mean +/− SEM. Statistical significance was calculated by one-way ANOVA *p< 0.05. N=number of mice, MNC=mono-nuclear cells as determined by forward/side scatter. For single knockout control mice, *Puma^−/−^*(N=7), *p21^−/−^*(N=8), *Zmat3^−/−^* (N=6), refer to Figure S1.

Previous reports have demonstrated important individual and overlapping roles of ZMAT3, PUMA and p21 in lymphoid cells (Deng *et al*, 1995; Villunger *et al*, 2003; Janic *et al*, 2018). Therefore, we hypothesised that *Puma^−/−^p21^−/−^Zmat3^−/−^* TKO mice may display a marked pre-leukaemic phenotype with a pronounced expansion of certain haematopoietic cell populations. We therefore determined the overall composition of the haematopoietic system of the TKO, and both DKO mice by immunostaining and fluorescence-activated cell sorter (FACS) analysis at 8-12 weeks of age. Cellularities of the different lymphoid organs (e.g. spleen, bone marrow, thymus) in the DKO and TKO mice were comparable to those in wt controls (Figures 1B, S1A), as were the white blood cell (WBC) counts in peripheral blood (PB) (Figure S1B). T cell development in the thymus appeared normal with the overall distribution of the double negative (DN, CD4^−^CD8^−^ progenitors), double-positive (DP, CD4^+^CD8^+^ immature) and single positive CD4^+^ and CD8^+^ (mature) thymocytes comparable between the DKO as well as TKO mice with those seen in wt controls (Figure S1C-D). In the bone marrow, we observed a small but significant increase in LSKs (Lineage marker– SCA1^+^cKIT^+^ HSPCs) in the *Puma^−/−^p21^−/−^Zmat3^−/−^* TKO mice compared to wt controls (Figure 1C, D). However, this did not result in any major differences in myeloid, erythroid or B cell development in the bone marrow (Figure 1E, S1E). The numbers of all mature cell types in the spleen that we tested were comparable between DKO as well as TKO mice with those seen in wt controls, with comparable frequencies of B cells (B220^+^), T cells (TCRβ^+^) and myeloid cells (MAC1^+^) (Figure 1F-H, S1G-I). In addition, histological analysis of major organs revealed no lesions of significance in young DKO and TKO mice (Figure S2). Overall, these findings demonstrate that the combined absence of ZMAT3, PUMA and p21 does not cause any marked defects in young pre-cancerous mice.

### Cells from *Puma^−/−^p21^−/−^Zmat3^−/−^* mice are resistant to TRP53-mediated apoptosis triggered by DNA damage

The BH3-only protein PUMA is critical for apoptosis induced by stress stimuli that activate TRP53, such as DNA damage (Villunger *et al*, 2003; Jeffers *et al*, 2003; Michalak *et al*, 2008). To verify that cells from *Puma^−/−^p21^−/−^Zmat3^−/−^* mice were, as expected, resistant to TRP53-mediated apoptosis, we exposed thymocytes from *Puma^−/−^p21^−/−^Zmat3^−/−^* as well as *Puma^−/−^Zmat3^−/−^, p21^−/−^Zmat3^−/−^*, *Zmat3^−/−^*, *p21^−/−^*, *Puma^−/−^* and wt mice to apoptosis inducing agents *in vitro* and their survival was then measured by flow cytometric analysis. The survival of *Puma^−/−^p21^−/−^Zmat3^−/−^* TKO thymocytes was comparable to that of DKO and wt thymocytes when either left untreated (DMSO control), treated with ionomycin or starved of serum (growth factor deprivation) (Figures 2A-C). As expected, thymocytes from *Puma^−/−^p21^−/−^Zmat3^−/−^* TKO and *Puma^−/−^Zmat3^−/−^* DKO mice were resistant to etoposide, a cytotoxic stimulus that kills these cells via a TRP53-dependent process. At 24 h, there was less than 40% survival of wt thymocytes but more than 80% survival of the *Puma^−/−^p21^−/−^Zmat3^−/−^* as well as *Puma^−/−^Zmat3^−/−^* and *Puma^−/−^* thymocytes (Figure 2D). These data show that cells from *Puma^−/−^p21^−/−^Zmat3^−/−^* mice are resistant to TRP53-dependent apoptosis triggered by DNA damage but normally sensitive to several TRP53-independent apoptotic stimuli.

**Figure 2.**
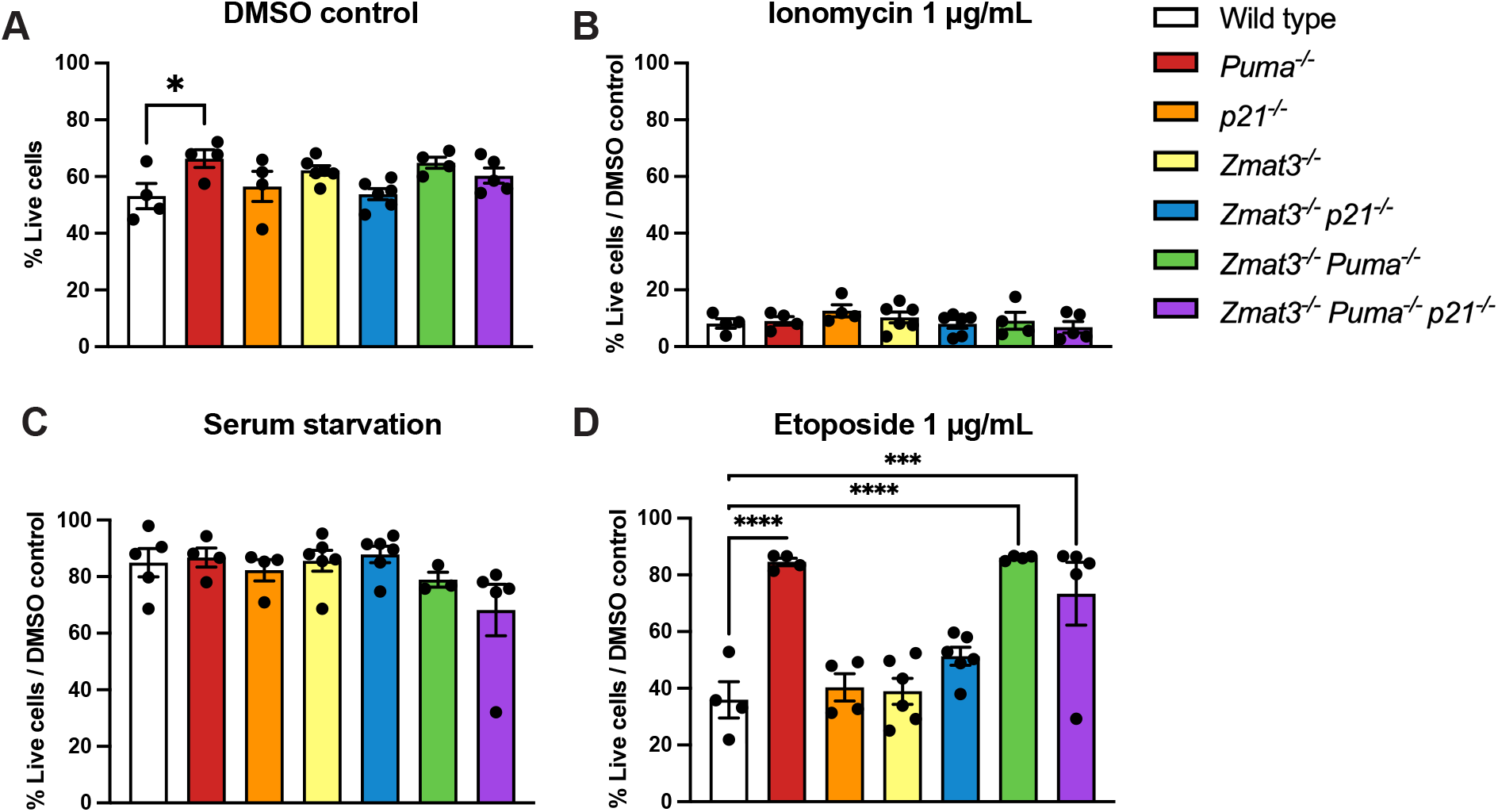
Impact of combined loss of ZMAT3, PUMA and p21 on thymocyte survival *in vitro* upon treatment with various apoptotic stimuli. Thymocytes from mice of the indicated genotypes were isolated and either **A)** left untreated (DMSO control), **B)** treated with 1 μg/mL ionomycin, **C)** exposed to serum deprivation in culture or **D)** treated with 1 μg/mL etoposide. Cell viability was assessed after 24 h by Annexin-V/PI staining and FACS analysis. Data are presented as mean +/− SEM of Annexin-V-PI-population (live cells) **(A)**, and live cells relative to DMSO treated control samples **(B-D)**. Statistical significance was calculated by one-way ANOVA *p= 0.035, ***p= 0.0009, ****p< 0.0001. N=number of mice of a particular genotype examined.

### Consequences of combined loss of ZMAT3 and p21 on γ-radiation induced thymic lymphoma development

Exposure of mice from a young age (starting at ~4 weeks) to repeated low-dose γ-radiation drives thymic lymphoma development through a mechanism that is suppressed by TRP53 (Kemp *et al*, 1994; Michalak *et al*, 2010). To investigate whether TRP53-mediated induction of ZMAT3 could restrain γ-radiation induced thymic lymphoma development, we examined the consequences of individual or combined loss of ZMAT3 and p21 in this model. Mice lacking PUMA alone or in combination with ZMAT3 and p21 were not included as it has been previously shown that mice defective in TRP53-induced apoptosis due to loss of PUMA are profoundly resistant to γ-radiation-induced thymic lymphoma development (Labi *et al*, 2010; Michalak *et al*, 2010). We found that *p21^−/−^Zmat3^−/−^* as well as *Zmat3^−/−^* and *p21^−/−^* mice developed thymic lymphoma at a similar rate to wt mice (Figure 3A). Immunophenotyping of the tumours from sick *p21^−/−^Zmat3^−/−^*, *Zmat3^−/−^* and *p21^−/−^* mice confirmed that these lymphomas were all of T lymphoid origin (Figure 3B), although lymphoma burden, as determined by thymus weight, was smaller in *p21^−/−^Zmat3^−/−^* mice compared to those detected in *Zmat3^−/−^*, *p21^−/−^* and wt mice (Figure 3C). Interestingly, high levels of p19ARF which are indicative of loss of TRP53 pathway function, were apparent in 3 out of 8 wt, 1 out of 3 *Zmat3^−/−^*, 1 out of 2 *p21^−/−^* but none out of 7 *p21^−/−^Zmat3^−/−^* lymphomas tested (Figures 3D-E). These findings demonstrate that in this mouse model of lymphomagenesis tumour suppression by TRP53 does not depend on ZMAT3 and/or p21. However, the absence of readily detectable defects in TRP53 pathway function in the lymphomas from the *p21^−/−^ Zmat3^−/−^* mice indicates that the absence of these two TRP53 target genes may obviate the selection for mutations in *Trp53* in this malignant disease.

**Figure 3.**
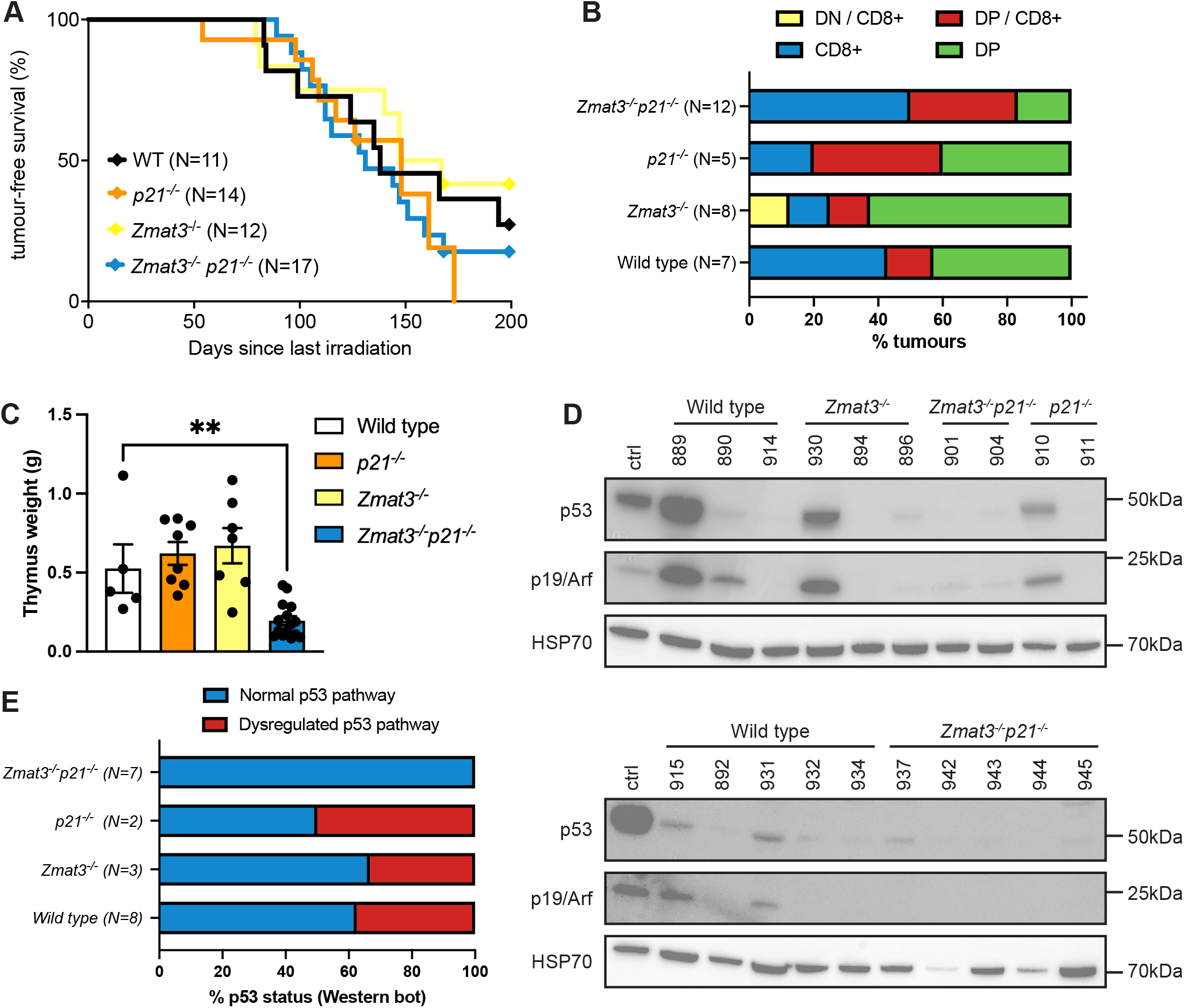
The combined absence of ZMAT3 and p21 does not accelerate γ-radiation induced thymic lymphoma development. **A)** Kaplan-Meier curves showing percentages of tumour-free mice of the indicated genotypes after exposure to four weekly doses of γ-radiation (1.5 Gy for each dose). Differences in thymic lymphoma incidence between wt and *p21^−/−^Zmat3^−/−^* were not statistically significant. P value determined by log-rank (Mantel-Cox) test p=0.5. **B)** Immunophenotyping of γ- radiation-induced thymic lymphomas arising in mice of the indicated genotypes, as assessed by cell surface marker staining and flow cytometric analysis of tumour cells from the thymus. Data are presented as the frequency of the indicated phenotypes for each genotype. Double negative CD4^−^CD8^−^ (DN) or double positive CD4^+^CD8^+^ (DP) lymphoma cells found in the thymus that show a trend towards the CD8 T cell lineage (DP/CD8^+^ or DN CD8^+^, respectively). N=number of mice. **C)** Thymus weights from sick mice of the indicated genotypes. Significant differences were observed in spleen weights between the sick *p21^−/−^ Zmat3^−/−^* and sick wt γ-irradiated mice. *p21^−/−^Zmat3^−/−^* (N=12), *p21^−/−^*(N=7), *Zmat3^−/−^*(N=7) and wt (N=5). Mean ± SEM, Unpaired Students t-test **p=0.0065. **D)** Western blot analysis of p19ARF, TRP53 and HSP70 (loading control) in thymic lymphomas of the indicated genotypes. The *Trp53* mutant *Eμ-Myc* lymphoma cell line EMRK1172 (Aubrey *et al*, 2015) was used as a control for p19ARF and mutant TRP53 protein overexpression. Protein size standards in kilodaltons are indicated. **E)** Percentages of the TRP53 status in γ-radiation-induced thymic lymphomas arising in mice of the indicated genotypes.

### *Zmat3^−/−^Puma^−/−^p21^−/−^* mice are prone to spontaneous tumour development

To examine the impact of combined loss of ZMAT3, PUMA and p21 in all tissues on the development of cancer, we aged *Puma^−/−^p21^−/−^Zmat3^−/−^* as well as *Puma^−/−^Zmat3^−/−^*, *p21^−/−^Zmat3^−/−^* and wt mice for 450-500 days. These animals were monitored for tumour development and major tissues were collected either at the time of sickness or at 450-500 days (study endpoint) regardless of health status. Interestingly, *Puma^−/−^p21^−/−^Zmat3^−/−^* as well as *Puma^−/−^Zmat3^−/−^* and *p21^−/−^Zmat3^−/−^* mice were significantly more prone to spontaneous tumour development compared to wt controls, with *Puma^−/−^p21^−/−^Zmat3^−/−^* mice showing a cancer incidence of nearly 50% by 500 days (Figure 4A). The majority of sick *Puma^−/−^Zmat3^−/−^* and *Puma^−/−^p21^−/−^ Zmat3^−/−^* mice presented with lymphoma (Figure 4B, Figure S3, Table 1) as determined by histopathological analysis, while most sick *Zmat3^−/−^p21^−/−^* mice presented with solid tumours (Table 1). None of the control w mice developed a tumour during the 500-day observation period (Figure 4A, B). Collectively, these results show that the combined loss of the three TRP53 target genes *Zmat3*, *Puma* and *p21* causes a high incidence of spontaneous tumour development. They also show that most malignancies arising in the compound mutant mice lacking PUMA are lymphomas whereas mice lacking ZMAT3 and p21 develop solid cancers.

**Table 1.**
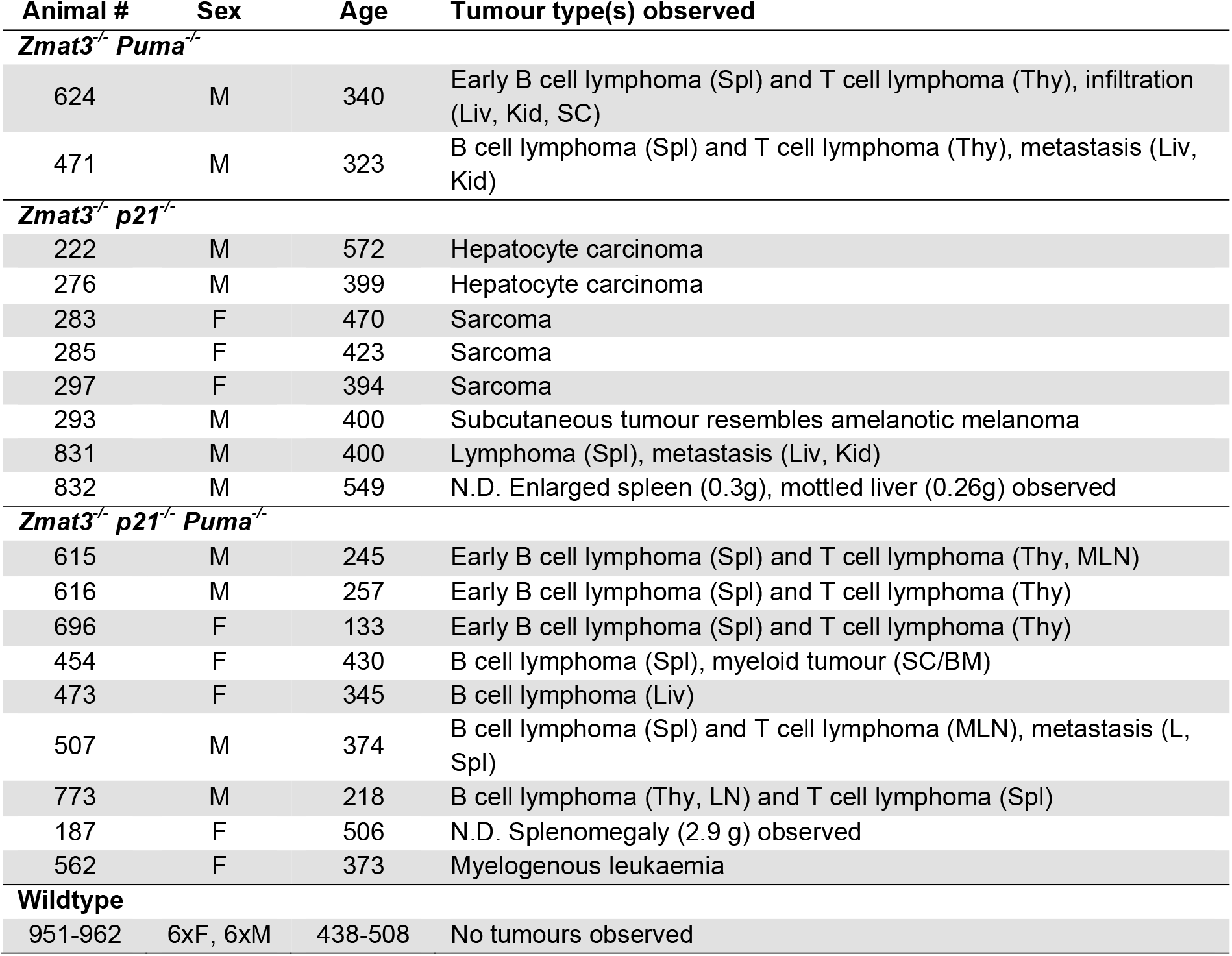
Tumour types observed in *Zmat3*, *Puma* and *p21* compound mutant mice. Aged mice from *Zmat3^−/−^ p21^−/−^ Puma^−/−^* inter-crosses were euthanised at ethical endpoint (sick, hind leg paralysis, ur volume, anaemia). Wild-type control animals were all healthy at time of analysis. Tumour types were mined by pathological analysis of H&E-stained organs and tumour sections. Organs involved noted in brackets; spleen; Thy, thymus; Liv, liver; Kid, kidney; SC, spinal cord; BM, bone marrow; MLN, mesenteric lymph node; L, LN, lymph node. N.D., histology not done, observations made after cryopreservation.

| **Animal #** | **Sex** | **Age** | **Tumour type(s) observed** |
| --- | --- | --- | --- |
| ***Zmat3*<sup>-/-</sup> *Puma*<sup>-/-</sup>** |  |  |  |
| 624 | M | 340 | Early B cell lymphoma (Spl) and T cell lymphoma (Thy), infiltration (Liv, Kid, SC) |
| 471 | M | 323 | B cell lymphoma (Spl) and T cell lymphoma (Thy), metastasis (Liv, Kid) |
| ***Zmat3*<sup>-/-</sup> *p21*<sup>-/-</sup>** |  |  |  |
| 222 | M | 572 | Hepatocyte carcinoma |
| 276 | M | 399 | Hepatocyte carcinoma |
| 283 | F | 470 | Sarcoma |
| 285 | F | 423 | Sarcoma |
| 297 | F | 394 | Sarcoma |
| 293 | M | 400 | Subcutaneous tumour resembles amelanotic melanoma |
| 831 | M | 400 | Lymphoma (Spl), metastasis (Liv, Kid) |
| 832 | M | 549 | N.D. Enlarged spleen (0.3g), mottled liver (0.26g) observed |
| ***Zmat3*<sup>-/-</sup> *p21*<sup>-/-</sup> *Puma*<sup>-/-</sup>** |  |  |  |
| 615 | M | 245 | Early B cell lymphoma (Spl) and T cell lymphoma (Thy, MLN) |
| 616 | M | 257 | Early B cell lymphoma (Spl) and T cell lymphoma (Thy) |
| 696 | F | 133 | Early B cell lymphoma (Spl) and T cell lymphoma (Thy) |
| 454 | F | 430 | B cell lymphoma (Spl), myeloid tumour (SC/BM) |
| 473 | F | 345 | B cell lymphoma (Liv) |
| 507 | M | 374 | B cell lymphoma (Spl) and T cell lymphoma (MLN), metastasis (L, Spl) |
| 773 | M | 218 | B cell lymphoma (Thy, LN) and T cell lymphoma (Spl) |
| 187 | F | 506 | N.D. Splenomegaly (2.9 g) observed |
| 562 | F | 373 | Myelogenous leukaemia |
| **Wildtype** |  |  |  |
| 951-962 | 6xF, 6xM | 438-508 | No tumours observed |

**Figure 4.**
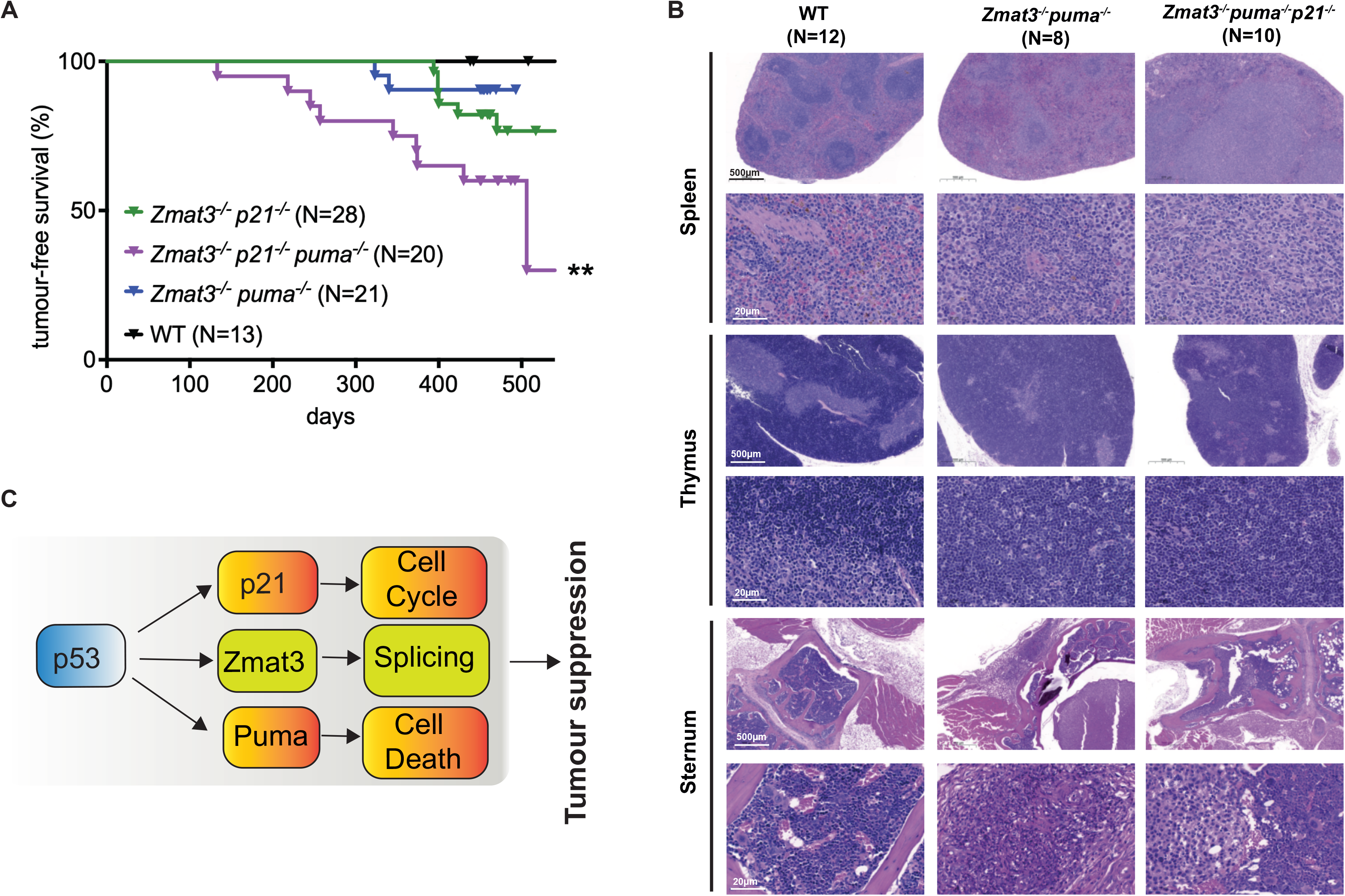
The combined absence of ZMAT3, PUMA and p21 causes spontaneous tumour development in mice. **A)** Kaplan-Meier curves showing tumour-free survival of *Puma^−/−^p21^−/−^Zmat3^−/−^*, *Puma^−/−^ Zmat3^−/−^, p21^−/−^Zmat3^−/−^* and control wt mice. Log-rank (Mantel-Cox) test P values comparing to wt mice; *Puma^−/−^p21^−/−^Zmat3^−/−^* p=0.003**; *Puma^−/−^p21^−/−^* p=0.06; *Puma^−/−^Zmat3^−/−^* p=0.26. **B)** Representative H&E-stained sections of spleen, bone marrow (sternum) and thymus of sick *Puma^−/−^p21^−/−^Zmat3^−/−^* (#454,615) and *Puma^−/−^Zmat3^−/−^* (#471,624) mice, and age-matched healthy wt (#953) mice for comparison. Scale bar denotes 500 μm or 20 μm, respectively. N=number of mice of a particular genotype analysed. **C)** Model: ZMAT3 loss drives spontaneous tumour development in collaboration with additional loss of PUMA and p21.

## DISCUSSION

TP53 is a critical suppressor of cancer development and progression (Kastenhuber & Lowe, 2017; Vousden & Lane, 2007; Thomas *et al*, 2022; Aubrey *et al*, 2018). TP53 functions as a transcription factor that can directly regulate the expression of ~500 target genes, and indirectly many more (Thomas *et al*, 2022; Boutelle & Attardi, 2021; Mello & Attardi, 2018). Transcriptional targets of TP53/TRP53 that are critical for the induction of apoptotic cell death, cell cycle arrest and senescence have been considered as essential effectors of TP53/TRP53-mediated tumour suppression (Boutelle & Attardi, 2021). Unexpectedly, however, genetic inactivation of all these TP53/TRP53-activated cellular processes does not render mice tumour prone (Valente *et al*, 2013; Brady *et al*, 2011a; Li *et al*, 2012). In contrast, TRP53-deficient mice spontaneously develop tumours with 100% of animals succumbing to malignant disease before 270 days of age (Donehower *et al*, 1992; Jacks *et al*, 1994). Genetic screens and genetically modified mouse strains have provided useful tools to examine the importance and biological functions of numerous TRP53 target genes in TRP53-mediated tumour suppression. Utilising *in vivo* CRISPR and shRNA screening in oncogene expressing mouse embryonic fibroblast *in vitro* (Bieging-Rolett *et al*, 2020) and haematopoietic stem/progenitor cells *in vivo* (Janic *et al*, 2018), respectively, the zinc finger RNA-binding protein ZMAT3 was identified as a critical factor contributing to TRP53-mediated tumour suppression.

Here we show that the combined absence of ZMAT3, PUMA and p21, the latter two critical for TRP53-mediated induction of apoptosis (Villunger *et al*, 2003; Jeffers *et al*, 2003) and cell-cycle arrest/cell senescence (Deng *et al*, 1995), respectively, resulted in nearly 50% of spontaneous tumour development by 500 days of age. Given that PUMA is a key factor in TRP53-mediated apoptosis in haematopoietic cells (Michalak *et al*, 2008; Jeffers *et al*, 2003; Villunger *et al*, 2003; Aubrey *et al*, 2018), it is perhaps not surprising that the majority of sick *Puma^−/−^Zmat3^−/−^* and *Puma^−/−^p21^−/−^Zmat3^−/−^* mice presented with lymphoma. Interestingly, some *Zmat3^−/−^p21^−/−^* sick mice developed a range of solid cancers at a relatively late age. Their incidence of such malignant disease was significantly higher compared to wt mice. This indicates that loss of apoptotic function, cell cycle arrest/cell senescence and ZMAT3-regulated alternative splicing or not yet identified functions of ZMAT3 contribute to the tumour suppressive function of TRP53 in both haematological and solid cancers, with the combination of the genes lost determining cancer type. Notably, *Puma^−/−^Zmat3^−/−^* and *Puma^−/−^ p21^−/−^Zmat3^−/−^* presented with lymphoma, and *Zmat3^−/−^p21^−/^* mainly solid tumours (6/8). This indicates that the loss of PUMA is particularly critical for spontaneous development of haematopoietic malignancies, consistent with the profound impact of its absence on the apoptosis of lymphoid and myeloid cells (Michalak *et al*, 2008; Jeffers *et al*, 2003; Villunger *et al*, 2003).

Examination of the lymphoid tissues from young *Puma^−/−^Zmat3^−/−^*, *Zmat3^−/−^p21^−/−^Puma^−/−^*and *p21^−/−^Zmat3^−/−^* mice revealed that they were largely normal, although we did observe a significant, albeit minor, increase in LSK HSPCs in *Puma^−/−^p21^−/−^Zmat3^−/−^* mice compared to wt controls. This is consistent with earlier reports that the absence of TRP53 itself does not cause detectable abnormalities in the haematopoietic system of young healthy mice although it predisposes them to spontaneous lymphoma development (Michalak *et al*, 2008; Strasser *et al*, 1994, Donehower *et al*, 1992; Jacks *et al*, 1994).

Several direct TP53 target genes have been implicated in DNA damage-induced apoptosis signalling, but only the BH3-only proteins, PUMA and to lesser extent NOXA, have been shown to be essential for TRP53-induced apoptosis (Villunger *et al*, 2003; Jeffers *et al*, 2003; Michalak *et al*, 2008, 2009). As expected, thymocytes from *Puma^−/−^p21^−/−^Zmat3^−/−^* and *Puma^−/−^Zmat3^−/−^* mice displayed significant resistance to etoposide (comparable to thymocytes from *Puma^−/−^* mice) to etoposide, a cytotoxic agent that kills these cells via a TRP53-dependent pathway (Strasser *et al*, 1994; Lowe *et al*, 1993; Clarke *et al*, 1993). This indicates that the additional loss of ZMAT3, alone or with further loss of p21, does not increase the protection against TRP53-dependent or TRP53-independent apoptosis that is caused by the absence of PUMA. Consistent with previous studies (Janic *et al*, 2018), this demonstrates that ZMAT3 does not have a prominent role in apoptosis, at least in lymphoid cells.

Previous reports in mice have shown that thymic T cell lymphoma development induced by γ-radiation can be significantly accelerated by loss of *Trp53* (even loss of one allele of *Trp53*) (Kemp *et al*, 1994; Kaplan & Brown, 1952; Labi *et al*, 2010) which impairs DNA damage-induced apoptosis, cell cycle arrest/cell senescence and coordination of DNA repair, which constitute critical processes for TRP53-mediated tumour suppression (Kastenhuber & Lowe, 2017; Thomas *et al*, 2022; Aubrey *et al*, 2018; Vousden & Lane, 2007; Wang *et al*, 2022). Whole body γ-irradiation activates TRP53 and causes induction of its downstream target genes *p21*, *Puma* and *Zmat3* in thymocytes as well as fibroblasts (Janic *et al*, 2018; Best *et al*, 2020) and probably many (if not all) other cell types. Here we investigated the contribution of ZMAT3 and p21 to the suppression of DNA damage induced thymic T cell lymphoma development. Of note, it was previously shown that the absence of PUMA abrogates γ-radiation induced thymic T cell lymphoma development because in this cancer model mobilisation and massive proliferation of HSPCs as a result of the depletion of mature blood cells is essential for tumour development, and the γ-radiation induced depletion of mature leukocytes is prevented by the absence of PUMA (Labi *et al*, 2010; Michalak *et al*, 2010). Therefore, *Puma^−/−^p21^−/−^Zmat3^−/−^* and *Puma^−/−^Zmat3^−/−^* mice were not included in these studies. Our investigations show that the absence of ZMAT3, either alone or in combination with loss of p21, did not accelerate (or slow) γ-radiation induced thymic T cell lymphoma development, in contrast to the loss of even a single allele of *Trp53*. Of note, while many *Zmat3^−/−^* and *p21^−/−^* thymic T cell lymphomas had acquired defects in TRP53 function, as revealed by high levels of p19ARF, all *p21^−/−^Zmat3^−/−^* lymphomas tested had retained wt TRP53 function. These findings demonstrate that ZMAT3 loss is not essential for the suppression of γ-radiation induced thymic T cell lymphoma development, probably because TRP53 can activate many at least partially redundant tumour suppressive processes in response to DNA damage. Coordination of DNA repair to maintain genomic stability is probably particularly important in this model. The absence of defects in TRP53 function in all *p21^−/−^Zmat3^−/−^* lymphomas tested may indicate that the combined loss of ZMAT3 and p21 obviates the selection for mutations in *Trp53* in this malignant disease.

In conclusion, our results confirm and extend the notion that ZMAT3 is a direct TRP53 target gene that contributes to TRP53-mediated tumour suppression. It probably does this by regulating a large number of mRNAs that function in the TP53 network, thereby impacting, perhaps in a feed-forward activation loop, several TP53-activated cellular responses (Bieging-Rolett & Attardi, 2021, Vilborg *et al*, 2011). Of note, on its own loss of ZMAT3 does not cause cancer but its tumour suppressive function becomes apparent when TRP53-mediated apoptosis (loss of PUMA) and/ or cell cycle arrest/cell senescence (loss of p21) are concomitantly disabled. Similarly, the impact of loss of ZMAT3 in mouse models of lung and liver cancer was less pronounced than that caused by the absence of TRP53 itself (Bieging-Rolett *et al*, 2020; Best *et al*, 2020). Additional support for the idea that in addition to the absence of ZMAT3, additional TRP53-regulated processes must also be disabled for spontaneous tumour development comes from the observation that mutations in *ZMAT3* are not prevalent in human cancers compared to the very high frequency of *TP53* mutations (Bieging-Rolett et al., 2020; Bieging-Rolett & Attardi, 2021). Thus, our findings further cement the importance of the coordinated action of multiple TP53/TRP53 activated cellular responses for tumour suppression. This may have implications for developing novel strategies for treating cancer as simultaneous activation of several of these responses might be needed for effective therapy.

## MATERIALS AND METHODS

### Mice

*Zmat3^−/−^* (Janic *et al*, 2018) and *Puma^−/−^p21^−/−^* (Valente *et al*, 2013) mice have been previously described. All mice were maintained on a C57BL/6-WEHI background. To produce *Puma^−/−^ p21^−/−^Zmat3^−/−^*, *Puma^−/−^Zmat3^−/−^* and *p21^−/−^Zmat3^−/−^* mice, we serially inter-crossed *Zmat3^−/−^* and *Puma^−/−^p21^−/−^* mice. All experiments with mice were performed with the approval of The Walter and Eliza Hall Institute Animal Ethics Committee and according to the Australian code of practice for the care and use of animals for scientific purposes. Genotyping was performed by polymerase chain reaction (PCR) to confirm the absence of *Zmat3, Puma* and *p21* (for PCR primers used, see Supplementary Table 1).

### Histology

Organs from mice were fixed in either 10% buffered formalin or Bouin’s solution and subsequently embedded in paraffin. Slides were prepared and stained with haematoxylin and eosin (H&E). Histological examination of the organs was performed by Phenomics Australia Histopathology and Slide Scanning Service, University of Melbourne.

### Thymic T cell lymphoma induction

*p21^−/−^Zmat3^−/−^*, *p21^−/−^, Zmat3^−/−^* and wt mice (starting at 27±6 days of age) were γ-irradiated weekly for 4 weeks with 1.5 Gy from a 60 Co source (Theratron Phoenix, Theratronics). Treated mice were then monitored for 200 days for signs of illness, and tumour onset was calculated from the last (4^th^) dose of γ-irradiation.

### Immunostaining and flow cytometric analysis

Thymus, spleen, and bone marrow were harvested, and single cell suspensions were prepared in PBS (Gibco) with 5 mM EDTA (Merck), supplemented with 5% foetal calf serum (FCS; Sigma-Aldrich) for staining. Antibody clones used are as follows; B220 (RA3-6B2), IgM (RMM-1), IgD (11.26C), TCRβ (H57.597), CD4 (GK1.5), CD8 (53-6.7), MAC1 (M1/70) SCA1 (E13-161.7), c-KIT (2B8), NK1.1(PK136), Ter119 (TER119), Ly6G (1A8-4-10-9), F4/80 (BM8), CD2 (RM2.1), and were obtained from eBioscience, BioLegend, or generated in-house. Staining with propidium iodide (PI Sigma-Aldrich; 1 μg/mL) was used to exclude dead cells. Whole-organ cell counts were determined by mixing a known concentration of APC Calibrite beads (Becton Dickinson) with each sample. Data were collected using LSRFortessa X-20 or FACSymphony analysers and examined using FlowJo 10 (Becton Dickson).

### Cell viability assays

Thymi were harvested from mice of the indicated genotypes and single cell suspensions were prepared by gentle mashing through a 100 μm nylon strainer (Falcon). Cells were cultured in high-glucose Dulbecco’s modified Eagle’s medium (DMEM) supplemented with 10% foetal bovine serum, 50 μM 2-mercaptoethanol, 100 mM asparagine, 100 U/mL penicillin, and 100 mg/mL streptomycin. 5×10^4^ cells were plated into 96-well flat-bottom plates in triplicate and treated, as described (Strasser *et al*, 1991), with DMSO (control), 1 nM dexamethasone (Sigma-Aldrich), 1 μg/mL etoposide (Sigma-Aldrich), 1 μg/mL ionomycin (Sigma-Aldrich) or subjected to serum starvation by culturing in medium containing only 1% foetal calf serum. Cell viability was assessed at 24 h by resuspending cells in Annexin V binding buffer (0.1 M Hepes (pH 7.4), 1.4 M NaCl, 25 mM CaCl_2_ containing fluorescein isothiocyanate (FITC)-conjugated Annexin-V and 1 μg/mL PI. Cells were examined by flow cytometric analysis using a BD-Biosciences LSR-II analyser. 10,000 events were recorded per sample and data were analysed using FlowJo 10 analysis software. Data are presented as the percentage of AnnexinV^−^PI^−^ (viable) cells relative to DMSO treated control samples. N = 4-5 mice/genotype.

### Western blot analysis

For Western blot analysis, cell lysates were prepared in radio-immunoprecipitation assay (RIPA) buffer supplemented with PhosSTOP and complete protease inhibitor cocktail (Roche). Protein concentration was determined by Bradford assay using the Protein Assay Dye Reagent Concentrate (Bio-Rad, Hercules, CA). Samples of 10 μg of protein were prepared in Laemmli buffer, boiled for 5 min and size-fractionated by gel electrophoresis on NuPAGE 10% Bis-Tris 1.5-mm gels (Life Technologies) in 2-(N-morpholino) ethanesulfonic acid (MES) buffer. Proteins were then transferred onto nitrocellulose membranes (Life Technologies) using the iBlot membrane transfer system (Bio-Rad). Antibody dilution and blocking were performed in 5% skim milk, 0.1% Tween 20 in phosphate-buffered saline (PBS). The following antibodies were used for probing; mouse TRP53 (clone CM5, Novocastra), p19/ARF (clone 5.C3.1, Rockland) and HSP70 (clone N6, gift from Dr. R Anderson, Olivia Newton-John Cancer Research Centre, Melbourne, VIC, Australia), the latter used as a control for protein loading. Secondary antibodies used include goat anti-mouse IgG and goat anti rabbit-IgG both conjugated to horseradish peroxidase (HRP) (Southern Biotech). Forte Western HRP substrate (Millipore, Billerica, MA) was used for developing the signal, and membranes were imaged and analysed using the ChemiDoc XRS1 machine with ImageLab software (Bio-Rad).

### ETHICS APPROVAL

All experiments with mice followed the guidelines of The Melbourne Directorate Animal Ethics Committee, according to The Walter and Eliza Hall Institute of Medical Research Ethics Committee.

## ACKNOWLEDGEMENTS

We thank all members of the Blood Cells and Blood Cancer Division at The Walter and Eliza Hall Institute (WEHI) for support and advice; G. Siciliano, D. Fayle, H. Johnson, C. Gatt, and their team for animal husbandry; Dr S. Monard and his team at the WEHI Flow Cytometry Unit for help with FACS analysis; Thomas Nikolaou and Kevin Weston for haematology support. This study utilised the Phenomics Australia Histopathology and Slide Scanning Service, University of Melbourne. This work was supported by grants and fellowships from the Australian Phenomics Network (APN); the Australian National Health and Medical Research Council (NHMRC) to M.J.H. and A.S. (1143105), Program Grant to A.S. (1016701), Fellowship to A.S. (1116937), Investigator grant to A.S. (2007887); the Leukemia and Lymphoma Society of America to A.S. and M.J.H. (LLS SCOR 7015-18); the Cancer Council of Victoria Project grant to A.S. (1052309) and Venture Grant to M.J.H. and A.S.; support to A.S. from the estate of Anthony (Toni) Redstone OAM and a Spanish Ministry of Economy and Development Grant to A.J. (RTI2018-099017-A-I00). A.J. is supported by Ramon y Cajal Research Fellowship (RYC2018-025244-I), A.S. and M.J.H. are supported by NHMRC Fellowships (1020363 and 1156095), M.S.B is supported by Cancer Council Victoria Postdoctoral Fellowship and Swedish Cancer Society (21 0355 PT). This work was made possible through the Victorian Government Operational Infrastructure Support and Australian Government, the “Unidad de Excelencia María de Maeztu’’ funded by the MCIN and AEI (CEX2018-000792-M) and La Caixa banking foundation (51110009).

## AUTHOR CONTRIBUTIONS

MSB, MJH, AJ and AS conceptualised and planned the study; MSB, KB, LG, GH, MJH, AJ and AS performed experiments, analysed data and/or wrote the paper.

## Supplementary Material

**Supplementary Figure 1. Lymphoid organ analysis of young adult mice of the indicated genotypes.**

Single-cell suspensions were prepared from spleen, thymus, peripheral blood and bone marrow of *Puma^−/−^p21^−/−^Zmat3^−/−^* (N=5), *Puma^−/−^Zmat3^−/−^* (N=2-4), *p21^−/−^Zmat3^−/−^* (N=6), *Puma^−/−^*(N=7), *p21^−/−^*(N=8), *Zmat3^−/−^*(N=6) and wt (N=7) mice and the indicated haematopoietic cell subsets were examined by immunostaining and FACS analysis. **A)** Total cell counts for bone marrow (1 femur,) whole spleen and thymus in mice of the indicated genotypes. **B)** Total white blood cell (WBC) counts in peripheral blood in mice of the indicated genotypes. **C)** Representative FACS plot of cells from a wt mouse indicating the gating strategy to identify cell populations in the thymus. Immature double-negative thymocytes (DN progenitor; CD4^−^ CD8^−^), double-positive thymocytes (DP immature; CD4^+^CD8^+^) and the mature CD4^+^CD8^−^ and CD4^−^CD8^+^ single-positive T lymphoid cell populations. **D)** Percentages of the indicated cell subsets in the thymus of mice of the indicated genotypes. **E-G)** Complementary data for Figure 1D-E, H to include single knockout mouse controls. **H)** Representative FACS plots of cells from wt mice indicate gating strategy to identify cell populations in the peripheral blood. B cells (B220^+^), T cells (TCRβ^+^) and myeloid cells (MAC1^+^B220^−^TCRβ^−^). **I)** Percentages of the indicated cell subsets in peripheral blood of mice of the indicated genotypes.

**Supplementary Figure 2. Major organ analysis of young adult mice of the indicated genotypes.**

Representative H&E-stained sections of the indicated organs, as indicated, from 8- to 12-week old mice of the indicated genotypes. N represented number of mice analysed. Scale bar denotes 500 μm.

**Supplementary Figure 3. Histological analysis of sick *Puma^−/−^p21^−/−^Zmat3^−/−^* mice.**

Representative H&E-stained sections of the indicated organs that show evidence of tumours from sick *Puma^−/−^p21^−/−^Zmat3^−/−^* mice with indicated mouse numbers. Age-matched wt mice were used for comparison (for summary of the histopathology report see Table 1). Scale bar denotes 500 μm.

